# Multiomic Analysis Reveals Molecular Pathways Associated with Intestinal Aggregation of α-Synuclein

**DOI:** 10.1101/2025.08.27.672706

**Authors:** Julia M. Balsamo, Ying Yan, Dylan Thai, Stephanie M. Cologna, Elizabeth N. Bess

**Affiliations:** Department of Chemistry, University of California, Irvine, CA 92617, USA; Department of Chemistry, University of Illinois Chicago, Chicago, IL 60607, USA; Laboratory for Integrative Neuroscience, University of Illinois Chicago, Chicago, IL, 60607; Department of Molecular Biology and Biochemistry, University of California, Irvine, CA 92617, USA

**Keywords:** α-synuclein, Parkinson’s disease, enteroendocrine cells, dopamine, nitrite, ROS, gut microbiome

## Abstract

Aggregates of the protein α-synuclein may initially form in the gut before propagating to the brain in Parkinson’s disease. Indeed, our prior work supports that enteroendocrine cells, specialized intestinal epithelial cells, could play a key role in the development of this disease. Enteroendocrine cells natively express α-synuclein and synapse with enteric neurons as well as the vagus nerve. Severing the vagus nerve reduces the load of α-synuclein aggregates in the brain, suggesting that this nerve is a conduit for gut-to-brain spread. Enteroendocrine cells line the gut lumen, as such, they are in constant contact with metabolites of the gut microbiota. We previously found that when enteroendocrine cells are exposed to nitrite—a potent oxidant produced by gut bacterial *Enterobacteriaceae*—a biochemical pathway is initiated that results in α-synuclein aggregation. Here, we determined that dopamine production is critical to this mechanism of nitrite-induced α-synuclein aggregation. Using enteroendocrine cells, we modulated dopamine biosynthesis and profiled the cellular proteome and lipidome. Proteomic signatures in dopamine-free cells were distinctly different than in enteroendocrine cells, highlighting pathways relevant to intestinal development of Parkinson’s disease. Intriguingly, we observed that enteroendocrine cells maintain viability upon exposure to nitrite and in the presence of α-synuclein aggregates. This cellular robustness suggests that dopamine-producing enteroendocrine cells may be a reservoir of toxic α-synuclein aggregates, which can spread through a prion-like process. As a possible antidote, our findings show that benserazide—a chemical inhibitor of dopamine biosynthesis—limits formation of these aggregates in enteroendocrine cells. These studies lay a foundation for mechanistically informed therapeutic targets to prevent intestinal formation of α-synuclein aggregates before they spread to the brain.

**For Table of Contents Use Only:** 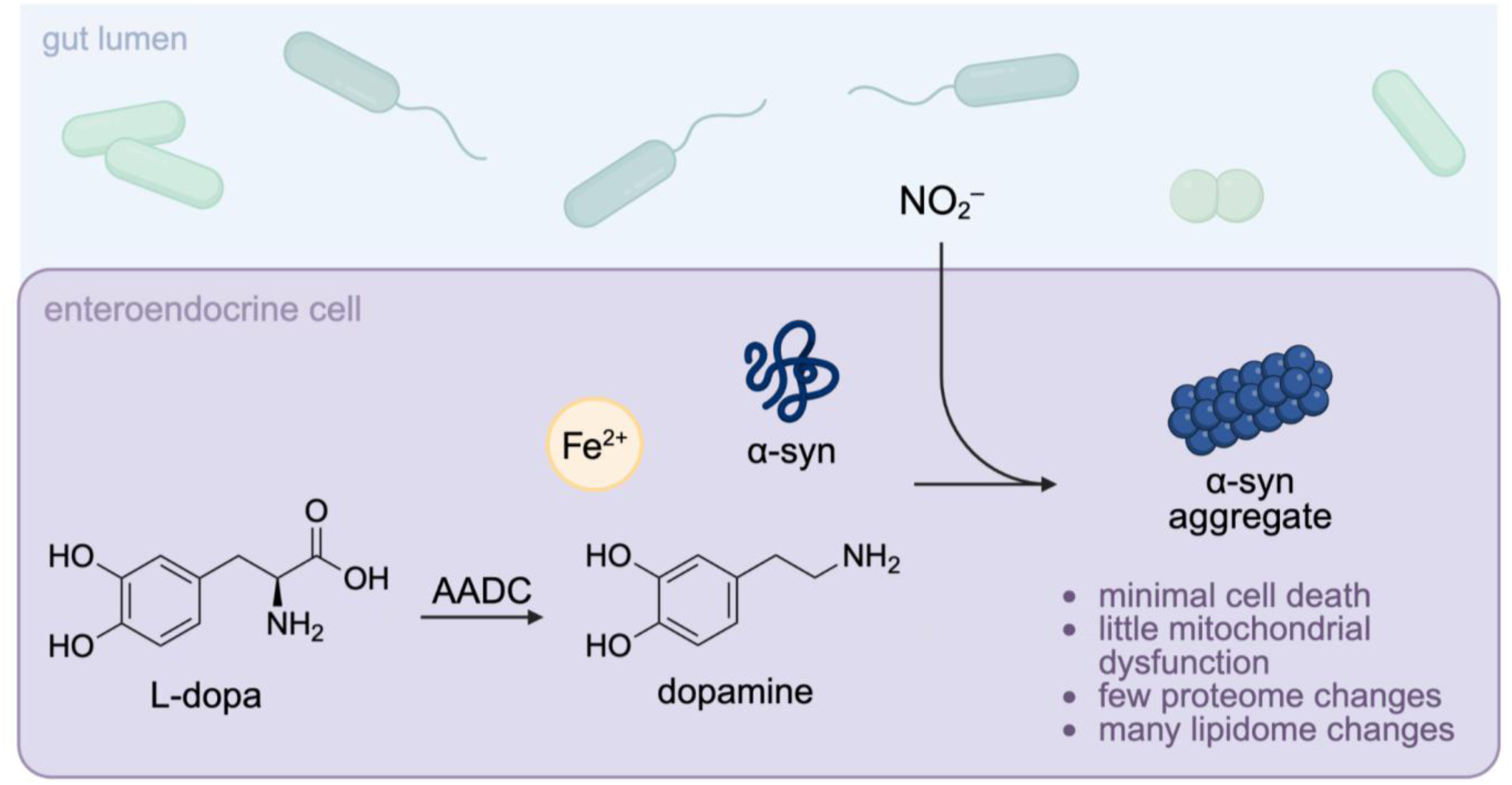

## INTRODUCTION

Parkinson’s disease (PD) has long been characterized by aggregation of the protein α-synuclein (α-syn) in dopaminergic neurons of the substantia nigra, a region of the brain that controls motor function.^1^ Our recent work suggests molecular-level mechanisms by which these toxic aggregates initially form in intestinal cells,^2^ from which they can propagate to the brain^3^ and advance degeneration of dopaminergic neurons over time.^4–6^ Enteroendocrine cells (EECs) located in the gut epithelium may be the source of intestinal α-syn, as they naturally express this protein.^7^ These cells also synapse with vagal neurons, which innervate the gut and brain.^8,9^ Through this connection, PD pathology may originate in the gut and spread to the brain via the vagus nerve;^10,11^ indeed, severing the murine vagus nerve inhibits gut-to-brain spread of α-syn.^12–14^

Only recently has the gut-specific etiology of PD begun to be explored.^15^ Accumulating evidence suggests a microbial component in the development of some PD subtypes, as the gut microbiome is significantly altered in people with PD as compared to non-diseased controls.^16–20^ Transplanting the gut microbiomes of people with PD into mouse models of this disease significantly exacerbated motor dysfunction as compared to the mice receiving a gut microbiome from people without PD.^21^ We previously reported a mechanism that may account for the gut microbiome’s role in PD;^2^ specifically, the role of *Enterobacteriaceae*—a family of bacteria that is enriched in the gut microbiotas of people with PD.^16,17,19^ *Enterobacteriaceae* produce the oxidizing agent nitrite upon nitrate respiration, a way that these bacteria harvest energy in the oxygen-limited intestinal lumen.^22^ We reported that nitrite instigates aggregation of natively expressed α-syn in EECs. Specifically, our complementary studies *in vitro* show that nitrite oxidizes Fe^2+^ to Fe^3+^, which subsequently oxidizes dopamine to dopamine-derived quinones that lead to aggregation of α-syn (Scheme 1).^2^ We also found that the extent of nitrite-induced α-syn aggregation depended on dopamine and its oxidation; strategies that limited dopamine oxidation mitigated aggregate formation.^23^

**Scheme 1.**
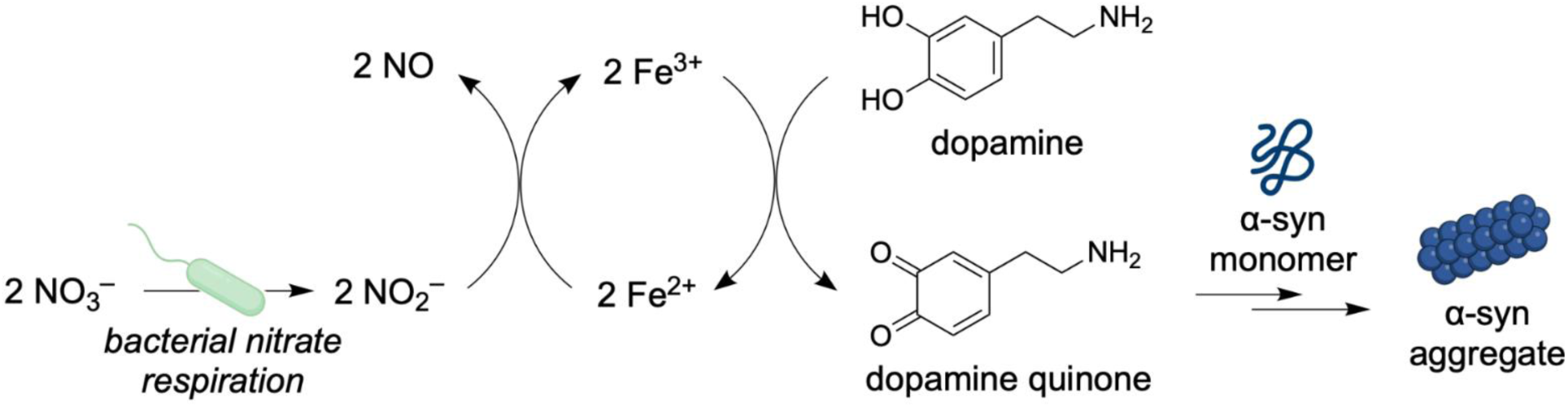
Nitrite-induced α-syn aggregation in enteroendocrine cells (EECs) bordering the gut lumen may depend on dopamine.

With our prior *in vitro* studies as our foundation, here we detail the mechanism of α-syn aggregation in intestinal EECs. We induce aggregation of native α-syn in model EECs using microbial metabolite nitrite. Dopamine’s role in nitrite-induced intestinal α-syn aggregation is deciphered using chemical tools to manipulate dopamine’s biosynthesis in EECs. Guided by studies of PD pathogenesis in brain-first subtypes of this disease, where cellular reactive oxygen species (ROS) are critical drivers of pathogenesis,^24–26^ we also characterize ROS generation and cell viability as a function of nitrite-induced α-syn aggregation. The cellular impacts of this process are also holistically profiled via proteomic and lipidomic profiling. Collectively, these studies provide a molecular detailing of intestinal α-syn aggregation and reveal new, needed possibilities for inhibiting α-syn aggregation in the intestine before it spreads to the brain.

## RESULTS

### Dopamine Biosynthesis Modulates α-Syn Aggregation in Gut Enteroendocrine Cells

We previously reported that nitrite induces α-syn aggregation in intestinal EECs (Scheme 1).^2^ Our findings from *in vitro* experiments show that dopamine and its subsequent oxidation, induced by nitrite, are critical to this pathogenic process.^23^ EECs naturally express α-syn as well as the dopamine metabolic pathway;^7,27^ here, we sought to delineate how intracellular levels of dopamine affect the ability of endogenous α-syn to aggregate in the context of the mammalian gut.

First, we investigated controlling the extent of α-syn aggregation by modulating dopamine biosynthesis. This biosynthesis begins with the amino acid tyrosine, which is converted to L-dopa by the enzyme tyrosine hydroxylase (Figure 1A).^28^ L-dopa is further converted to dopamine via aromatic amino acid decarboxylase.^29^ Benserazide (Benz) prevents production of dopamine from L-dopa by inhibiting aromatic amino acid decarboxylase.^30^ We hypothesized that supplying benserazide to EECs could reduce nitrite-induced dopamine-dependent α-syn aggregation by limiting endogenous dopamine levels.

**Figure 1.**
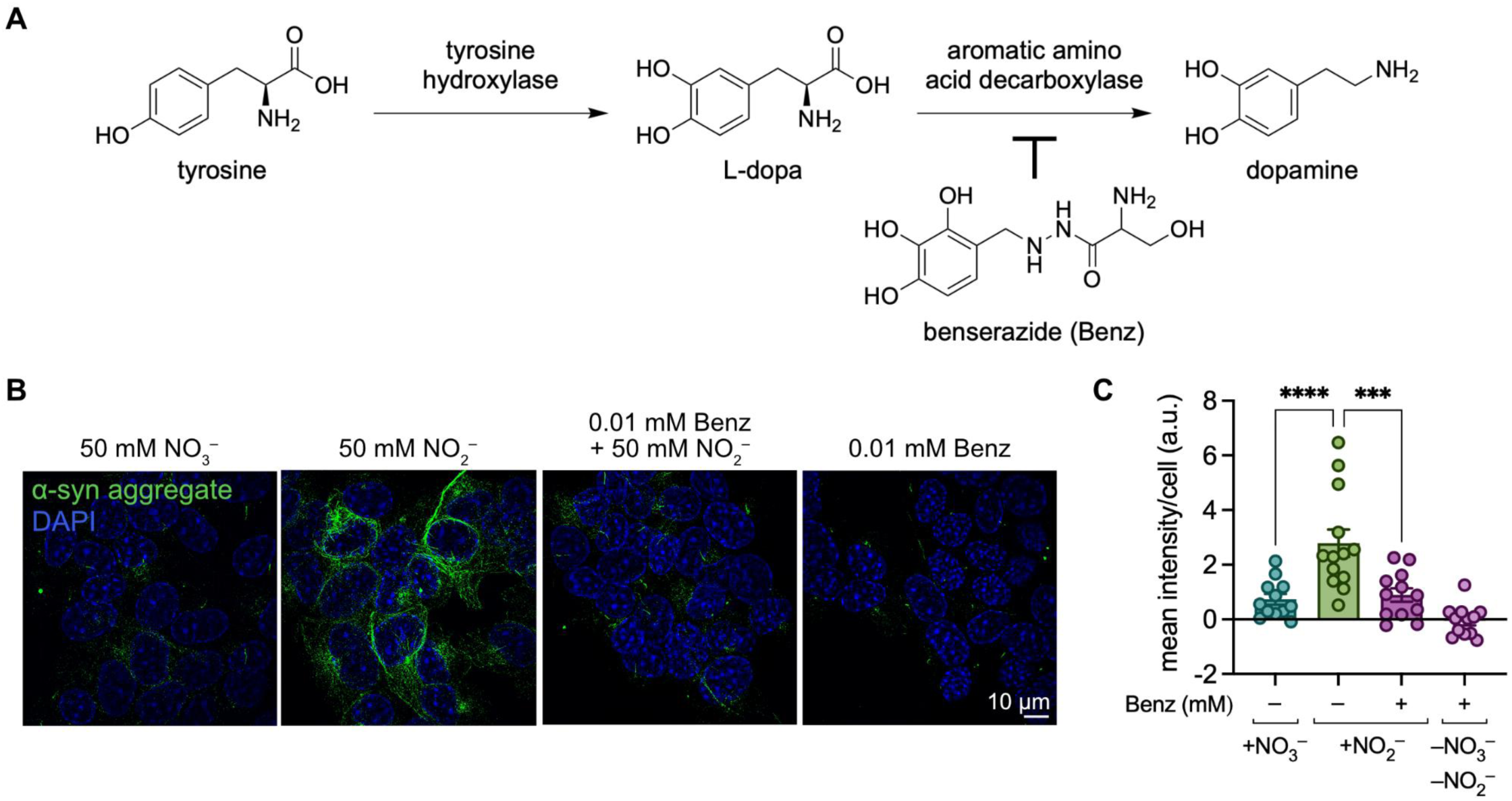
Nitrite-induced dopamine-dependent α-syn aggregation in enteroendocrine cells (EECs) is reduced in the presence of an inhibitor of dopamine biosynthesis. (A) Putative dopamine biosynthesis pathway within EECs. (B) Representative images of fixed STC-1 cells incubated for 24 h with 50 mM nitrate, 50 mM nitrite, and/or 0.01 mM benserazide (Benz), probed with anti-α-syn aggregate primary antibody MJFR-14. α-Syn aggregate signal is in green, and DAPI-stained nuclei are in blue. (C) Mean fluorescence intensity per cell was quantified from maximum intensity projections acquired by structured illumination microscopy (*n* = 3 independent biological replicates, 3–4 technical replicates for each, bars denote mean ± S.E.M.; significance determined by one-way ANOVA with Sidak’s multiple comparison test, ****: *P* < 0.0001, ***: *P* = 0.0002).

To investigate the impact of this enzyme inhibitor on α-syn aggregation, STC-1 cells (a murine cell line that is an accepted model system for EECs^31^) were incubated with nitrite (50 mM) with or without benserazide (0.01 mM) for 24 h. Benserazide-only-treated (0.01 mM) and untreated STC-1 cells were included as controls. Fixed STC-1 cells were probed using an antibody with an affinity for α-syn aggregates (MJFR-14)^2,32^ to allow for quantifying α-syn aggregation via immunofluorescence microscopy (Figure 1B). Isotype controls support no significant non-specific binding (Figure S1). Supplying benserazide to nitrite-treated cells attenuated α-syn aggregation by 3.1-fold relative to EECs treated solely with 50 mM nitrite (Figure 1C). This finding strongly supports that dopamine mediates α-syn aggregation in cells, analogous to our findings *in vitro*.^2,23^

### Mitochondria Remain Functional Overall in Nitrite-Treated Gut Enteroendocrine Cells

Mitochondrial dysfunction is thought to be a hallmark of PD;^24–26,33,34^ thus, we screened for mitochondrial dysfunction in EECs to further explore molecular pathways affected by intestinal aggregation of α-syn. To this end, STC-1 cells were probed with the fluorescent mitochondrial superoxide (MitoSOX) indicator, which enables quantification of mitochondrial dysfunction as a function of superoxide generation. We previously observed significant intracellular α-syn aggregation at concentrations as low as 0.5 mM nitrite;^2^ thus, we expected to detect significant ROS generation both at and above that concentration. To this end, we exposed STC-1 cells to nitrite and nitrate (0.005–50 mM) as well as antimycin A (AMA, 50 μM)—a known mitochondrial electron transport chain inhibitor commonly used as a positive control for superoxide generation.^35,36^ Surprisingly, across three independent studies (n = 9 total biological replicates), only at the highest concentration of nitrite supplied (50 mM) is there significant superoxide generation as compared to untreated cells (Figure 2A). These data indicate that no significant mitochondrial dysfunction occurs as a result of nitrite exposure except at relatively high concentrations. Additionally, despite nitrite being an oxidant, supplying it to EECs does not result in a cellular environment of oxidative stress at physiological concentrations of nitrite (0.5 mM)^37^ that induce α-syn aggregation. These results reveal that EECs can maintain mitochondrial function despite the presence of nitrite and toxic α-syn aggregates.

**Figure 2.**
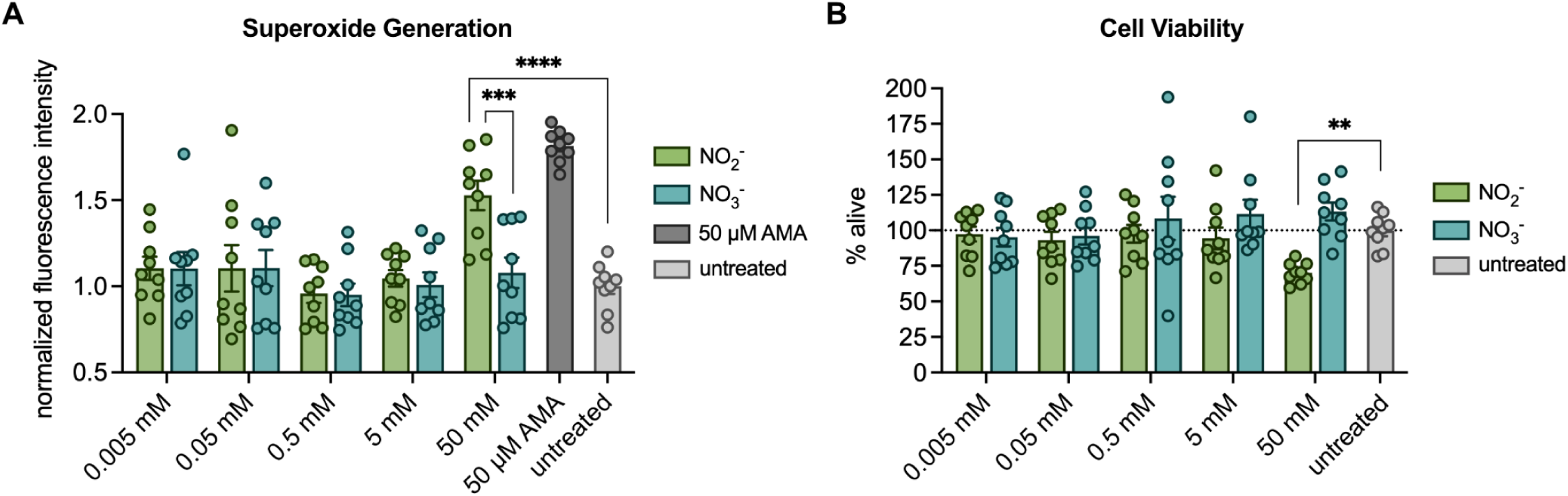
Gut enteroendocrine cells (EECs) have few biochemical defects despite the presence of the oxidant nitrite and toxic α-syn species. (A) Quantification of mitochondrial dysfunction as a function of superoxide generation in STC-1 cells after 24 h incubation with nitrite or nitrate (0.005–50 mM) using a fluorescent MitoSOX indicator. (B) Quantification of STC-1 cell viability after a 24 h incubation with nitrite or nitrate (0.005–50 mM), assessed via a colorimetric MTT assay. All error bars represent S.E.M.; n = 3 independent experiments, each with 3 biological replicates; significance determined by one-way ANOVA with Sidak’s multiple comparison test, ****: *P* < 0.0001, ***: *P* = 0.0003, **: *P* = 0.0064.

### Gut Enteroendocrine Cells Remain Viable Despite the Presence of Toxic α-Syn Species

PD is often characterized by dopaminergic neuronal cell death linked to α-syn aggregation in the substantia nigra.^38^ We sought to investigate the cytotoxicity of nitrite-induced α-syn aggregation in EECs. The viability of STC-1 cells treated with varying concentrations of nitrite or nitrate (0.005–50 mM) was determined by the colorimetric thiazolyl blue tetrazolium bromide (MTT) assay. In this assay, live cells convert MTT to formazan dye, which is quantified as a proxy for cell viability. Cytotoxicity relative to untreated cells was only observed at the highest concentration of nitrite tested (50 mM), where a 30% decrease in viability was detected; thus, nitrite’s impact on cell viability is minor (Figure 2B). It is noteworthy that the MTT assay relies on mitochondrial reductase enzymes to assess cytotoxicity. The minor extent of cytotoxicity observed at 50 mM nitrite thereby corroborates the data provided by the MitoSOX assay (mitochondrial dysfunction is present but only to a minor extent). Specifically, these data indicate that gut enteroendocrine cells are highly durable to nitrite and pathogenic α-syn aggregates, and defects are likely observed due to mitochondrial dysfunction. The absence of significant cell dysfunction or death could mean that nitrite-induced α-syn aggregates can persist within cells, increasing the likelihood of their characteristic prion-like spread and their subsequent role in PD pathogenesis.

### Nitrite-Induced α-Syn Aggregation in Gut Enteroendocrine Cells Minimally Activates the Proteome

Intrigued by the largely insignificant impacts of nitrite-induced α-syn aggregation on ROS generation and cell toxicity, we sought to comprehensively interrogate impacts of this process by profiling the cellular proteome. Aggregation of endogenous α-syn was first stimulated in STC-1 cells using nitrite (50 mM). Although higher than the physiological concentration in the intestinal lumen,^37^ lower concentrations of nitrite did not induce significant mitochondrial dysfunction or cytotoxicity (Figure 2). Cells were also treated with nitrate (50 mM) or vehicle, serving as negative controls.

To verify aggregation in the presence of nitrite and its absence in the nitrate and untreated cells, immunofluorescence staining using an antibody with affinity for α-syn aggregate was performed as previously described (vide supra). Upon verifying α-syn aggregation was induced in STC-1 cells upon nitrite treatment (Figure S2), the cells were subsequently lysed, and proteins were extracted and proteolytically cleaved. Tryptic peptides were separated by reversed-phase nanoflow liquid chromatography and analyzed by mass spectrometry. Protein identification and relative quantification was obtained and comparative analysis was performed to evaluate the differential proteome of the nitrite-treated, nitrate-treated, and untreated STC-1 cells. We observed that, of the 3708 identified proteins, 73 proteins were downregulated and 85 were upregulated in the nitrite-treated group compared to untreated cells (adjusted P < 0.05) (Figure 3A and Table S1). Meanwhile, of the 3704 proteins identified in the nitrate-treated group, 44 were downregulated and 103 were upregulated compared to untreated cells (Figure S3 and Table S1). When comparing the nitrite-treated cells to the nitrate-treated cells, we found that 111 proteins were downregulated and 49 were upregulated (Figure 3B and Table S1). Curiously, the concentration of total α-syn did not significantly change relative to untreated controls (adjusted P = 0.948), which suggests that despite aggregation, the expression level of α-syn does not significantly change.

**Figure 3.**
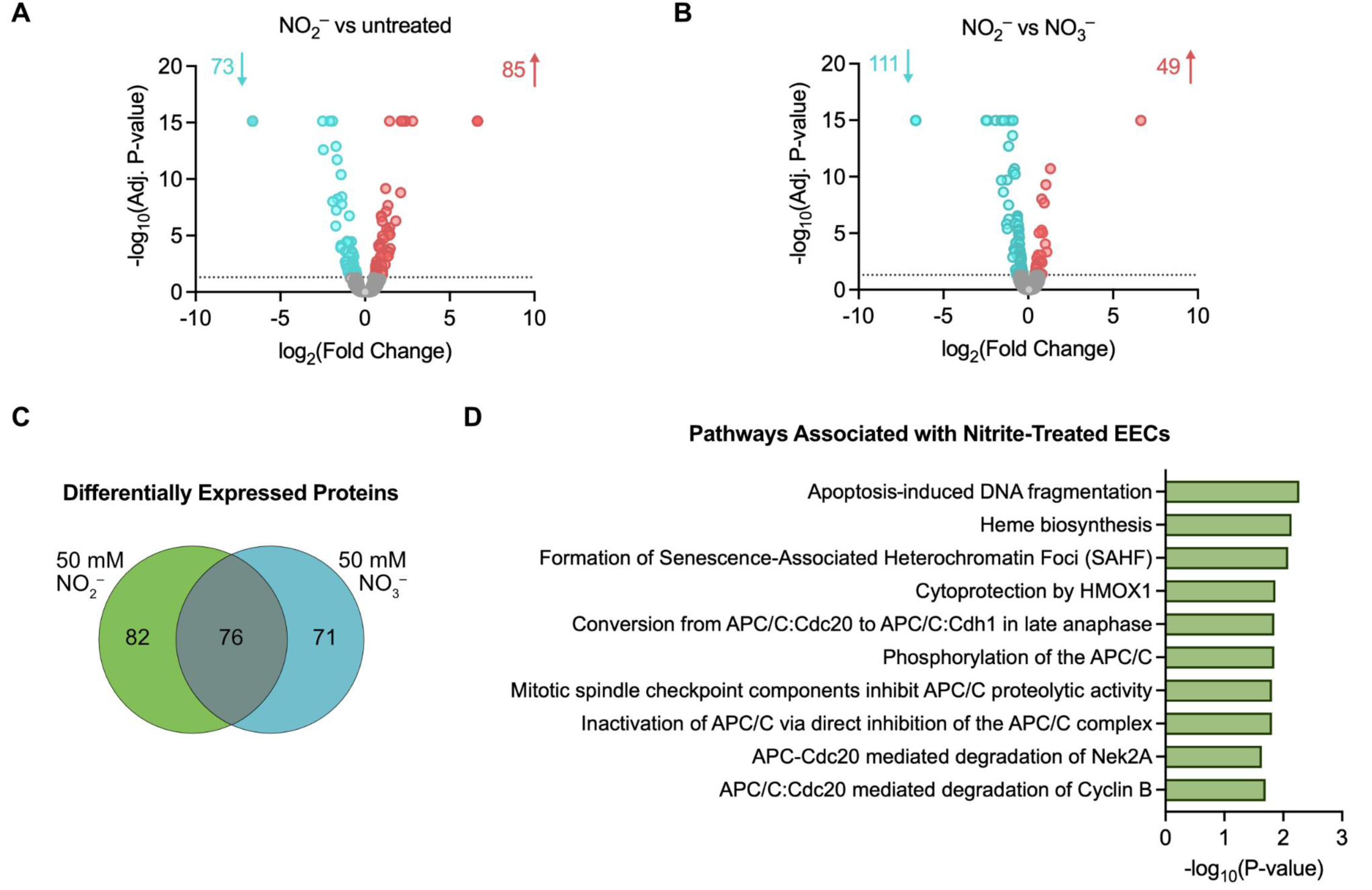
Differential proteomics of STC-1 enteroendocrine cells (EECs) exposed to nitrite—the product of bacterial nitrate respiration. (A–B) Volcano plots showing differential expressed proteins (cyan = downregulated, coral = upregulated) in enteroendocrine STC-1 cells treated with 50 mM nitrite, 50 mM nitrate, or vehicle. The P-value cutoff was 0.05. (A) In the nitrite group, 3708 proteins were identified; 73 were downregulated and 85 were upregulated as compared with the untreated group. (B) In the nitrite group, 3708 proteins were identified; 111 were downregulated and 49 were upregulated as compared with the nitrate-exposed group. (C) Venn diagram showing uniquely altered proteins in nitrite-treated cells as compared with the differential proteome of nitrite-treated relative to untreated cells and that of nitrate-treated relative to untreated cells (D) Pathway enrichment analysis of differential proteins of nitrite-treated STC-1 cells. Proteins unique to nitrite-treated STC-1 cells were identified after comparison between significantly altered proteins in both the nitrite-treated vs untreated cells and the nitrate-treated vs untreated cells.

When we compared the differential proteomes of nitrite-treated cells relative to untreated cells (158 proteins) and that of nitrate-treated cells relative to untreated cells (147 proteins), a total of 82 proteins were uniquely altered in nitrite-treated cells (Figure 3C and Table S2). Pathway analysis of the 82 uniquely differentially expressed proteins was performed using Reactome to elucidate the biological relevance of these proteomic shifts. The most significantly altered pathways are related to apoptosis-induced DNA fragmentation (P = 0.0054), heme biosynthesis (P = 0.00726), formation of senescence-associated heterochromatin foci (SAHF) (P = 0.00823), cytoprotection by heme oxygenase-1 (HMOX1) (P = 0.01362), and six pathways associated with the anaphase-promoting complex/cyclosome (APC/C) (Figure 3D). Apoptotic DNA fragmentation is a hallmark of programmed cell death that can be triggered by ROS.^39^ This finding aligns with our prior results: ROS was elevated in STC-1 cells treated with nitrite (50 mM) and corresponded with a decrease in cell viability (Figure 2A and 2B). Curiously, SAHF are structures that form in the nucleus when cells are no longer proliferating and have instead entered cellular senescence; this protective mechanism is stimulated by stressors, including oxidative stress and DNA damage.^40^ The presence of SAHF supports the idea of intestinal cells in a state of cell-cycle arrest despite continuously propagating α-syn aggregates.

The six pathways associated with the APC/C further elucidate the cellular response of EECs to the accumulation of toxic α-syn aggregates. The APC/C is an E3 ubiquitin ligase that tags proteins for degradation and plays a pivotal role in mitosis.^41^ Phosphorylation of the APC/C (P = 0.01420), inhibition of the APC/C proteolytic activity by mitotic spindle checkpoint components (P = 0.01557), and inactivation of the APC/C via direct inhibition of the APC/C (P = 0.01557) (Figure 3D) could be linked to both the accumulation of α-syn aggregates within nitrite-treated EECs and to the transition to cellular senescence to prevent further damage.

### Nitrite Exposure Modulates Different Protein Pathways in Dopamine-Free Cells

Toward disentangling the cellular impacts of nitrite versus those of α-syn aggregation on cellular processes, we turned to HeLa cells, which natively express α-syn but have no detectable dopamine in cell lysates.^42^ Exposing HeLa cells to nitrite (50 mM; 24 h) resulted in no significant increase in α-syn aggregation relative to cells treated with nitrate (Figure 4A and S4). The absence of nitrite-induced α-syn aggregation in dopamine-free HeLa cells is consistent with our findings using STC-1 cells, which show that dopamine biosynthesis is critical to the occurrence of nitrite-induced α-syn aggregation (Figures 1B and C).

**Figure 4.**
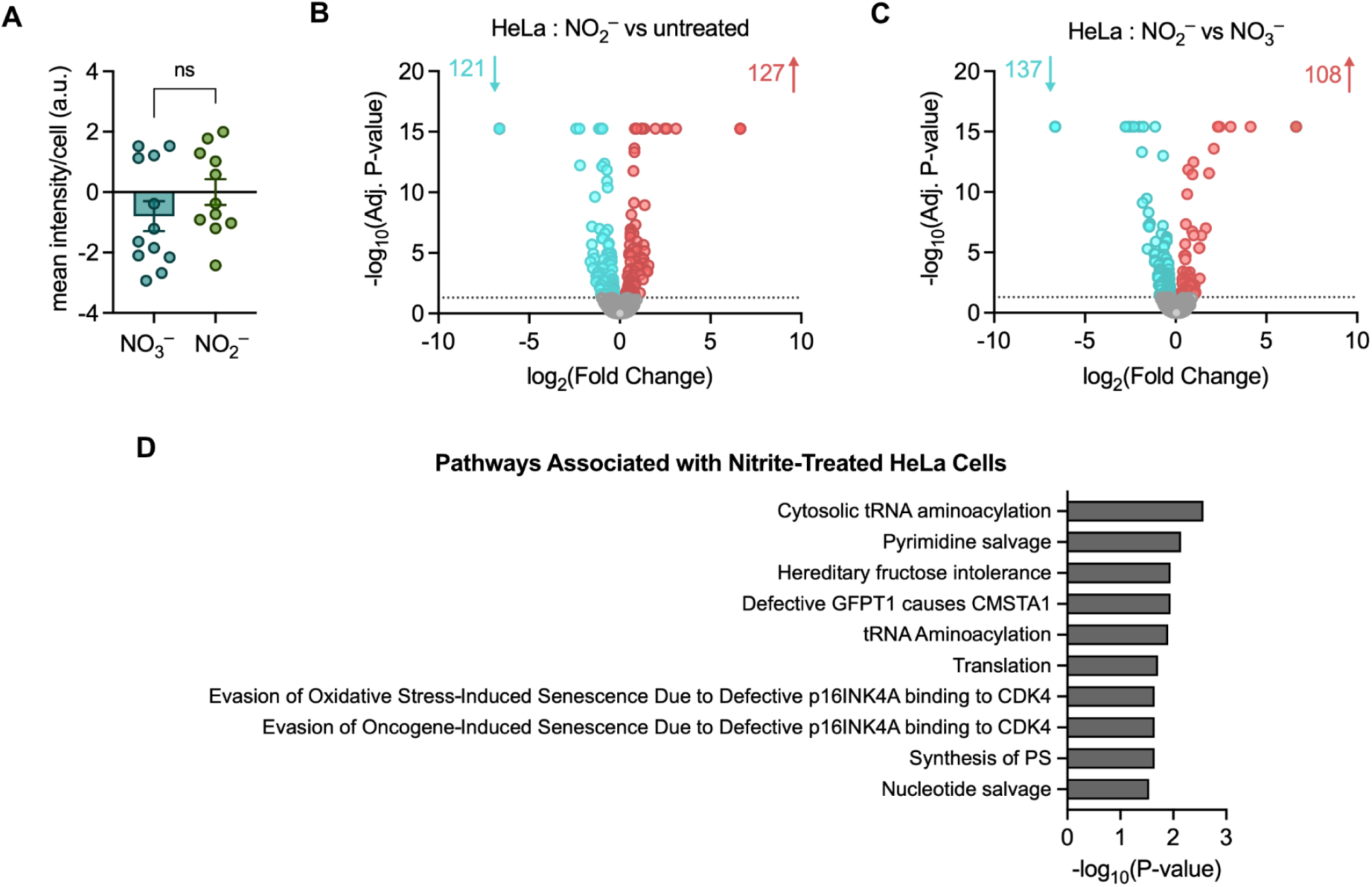
Differential proteomics of dopamine-free cells exposed to nitrite or nitrate. (A) HeLa cells were exposed to either 50 mM nitrite, 50 mM nitrate, or vehicle for 24 h. Mean fluorescence intensity per cell was quantified from maximum intensity projections acquired by structured illumination microscopy (*n* = 3 independent biological replicates, 3–4 technical replicates for each, untreated signal subtracted as background, bars denote mean ± S.E.M.; significance determined by t-test. (B–C) Volcano plots showing differential proteins (cyan = downregulated, coral = upregulated) of HeLa cells treated with 50 mM nitrite, 50 mM nitrate, or vehicle. The P-value cutoff was 0.05. (B) In the nitrite-treated group compared with the untreated HeLa cells, 4035 proteins were identified: 121 were downregulated and 127 were upregulated. (C) In the nitrite-treated group compared with nitrate-treated HeLa cells, 4034 proteins were identified: 137 were downregulated and 108 were upregulated. (D) Pathway enrichment analysis of unique differentially expressed proteins of the nitrite-treated HeLa cells. Proteins unique to nitrite-treated HeLa cells were identified after comparison between significantly altered proteins in the nitrite-treated vs untreated cells and the significantly altered proteins in the nitrate-treated vs untreated cells.

The absence of α-syn aggregation in HeLa cells provided an opportunity to distinguish the distinct impacts of nitrite and α-syn aggregation on cell function. HeLa cells were incubated with either nitrate (50 mM), nitrite (50 mM), or vehicle. Cell lysates were prepared as noted above for mass spectrometry-based proteomics. A total of 4035 proteins were identified when comparing the nitrite-treated and untreated samples. We observed that 121 proteins were downregulated and 127 proteins were upregulated relative to the untreated control (adjusted P < 0.05) (Figure 4B and Table S3). When comparing the nitrate-treated cells to the untreated cells, a total of 4031 proteins were identified, with 101 proteins downregulated and 137 proteins upregulated (Figure S5 and Table S3). When examining proteome changes in nitrite-treated cells compared with nitrate-treated cells, the nitrite-exposed cells displayed 137 downregulated proteins and 108 upregulated proteins (Figure 4C and Table S3).

Next, we sought to examine the uniquely altered proteins within the nitrite-treated cells. We compared the differential proteomes of nitrite-treated cells relative to untreated cells (248 proteins) and that of nitrate-treated cells relative to untreated cells (238 proteins) and found a total of 132 proteins were uniquely altered in nitrite-treated cells (Table S4). The 132 uniquely altered proteins identified were subjected to pathway enrichment analysis using Reactome and found to be most associated with cytosolic tRNA aminoacylation (P = 0.0027), pyrimidine salvage (P = 0.0071), and hereditary fructose intolerance (P = 0.0113) (Figure 4D). The top three most significantly altered pathways (P < 0.05) upon nitrite exposure include and are primarily associated with cellular, nucleotide, and energy metabolism. These stand in contrast to the significantly altered pathways observed in nitrite-treated STC-1 cells—specifically, those linked to apoptosis (apoptosis-induced DNA fragmentation), cellular aging and oxidative stress linked to neurodegenerative disease (SAHF), and ubiquitin E3 ligases involved in protein degradation (APC/C protein complex). In nitrite-treated HeLa cells, the lack of both α-syn aggregation and proteome alterations that we associated with nitrite-induced α-syn aggregation in STC-1 cells indicate that formation of α-syn aggregates, not exposure to oxidizing agent nitrite, is the driver of proteomic shifts observed in STC-1 cells. Additionally, the deficiency of dopamine in HeLa cells highlights the significance of endogenous dopamine and the relevance of STC-1 cells to intestinal α-syn aggregation.

### Nitrite Exposure Significantly Activates the Lipidome of Enteroendocrine Cells

Accumulating evidence has emphasized the role of lipid metabolism dysfunction in PD pathology, suggesting the disease is not only a disease characterized by misfolded proteins (a proteinopathy) but also a lipidopathy, a disease of disordered lipid metabolism.^43–45^ Indeed, studies using untargeted high-performance liquid chromatography-tandem mass spectrometry performed on human serum and cerebrospinal fluid from people with idiopathic and genetic PD found significant alterations in several lipid species as compared with non-diseased controls.^46^ The gut microbiota has since also been implicated in altered lipid and energy metabolism in PD.^47^ Bidirectional interactions between lipids and gut bacteria—the composition of which can significantly change depending on dietary lipid intake—have also been observed.^48^ This is particularly relevant for people taking a functional food approach in mitigating PD.^49–51^ We thus sought to characterize the lipidome of EECs upon nitrite-induced α-syn aggregation to better understand biologically relevant alterations of lipids in the gut.

Lipids were profiled in STC-1 cells treated with either nitrite (50 mM), nitrate (50 mM), or vehicle for 24 h. Lipid extracts were analyzed in both positive- and negative-ion modes via liquid chromatography–mass spectrometry (LC–MS) for untargeted lipidomics. A total of 282 distinct lipid species were identified by LC-MS. Members of many lipid classes were identified across treatment groups in EECs (Figure 5A and Table S5). The abundance of all lipids was profiled across the three experimental groups. Upon comparing nitrite-treated and untreated STC-1 cells, 90 lipids were found to be significantly more abundant in the nitrite group, while no lipids were found to be significantly reduced (P ≤ 0.05) (Figure S6A and Table S5). In the nitrate-treated group, 68 lipids were increased and 17 lipids were decreased as compared to untreated cells (Figure S6B and Table S5). Comparing nitrite- and nitrate-treated cells, one lipid was less abundant and 46 lipids were more abundant (Figure S6C and Table S5).

**Figure 5.**
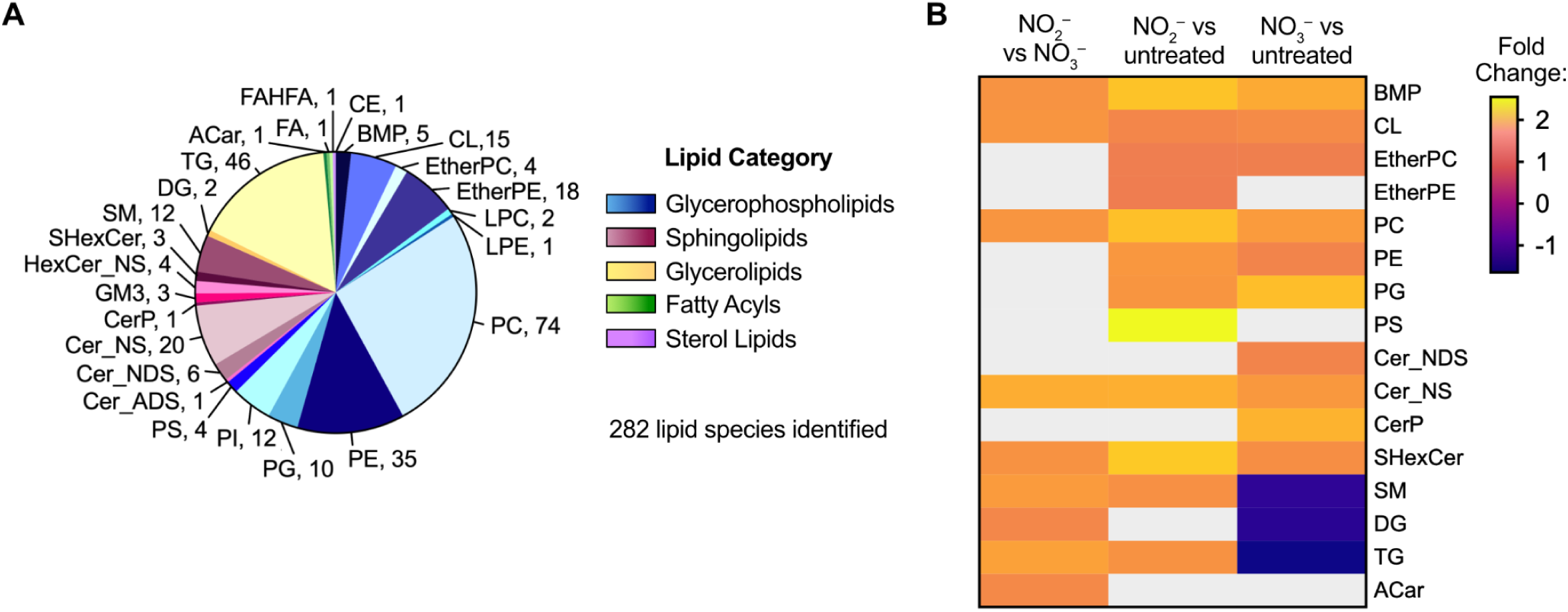
Untargeted lipidomics of STC-1 enteroendocrine cells (EECs) treated with nitrite. (A) All 282 lipids identified are shown, grouped by lipid class and category. (B) Class-based heat map depicting median fold-change of significantly altered lipid species across different treatment groups (50 mM nitrite, 50 mM nitrate, and untreated). Boxes in grey denote excluded lipid classes due to non-significant P-values; P-value cutoff was 0.05. Lipid abbreviations are denoted in Table S6.

Upon calculating the median fold-change for each class using significantly altered lipid species, a notable trend was visualized for the sphingolipid, sphingomyelin (SM) (Figure 5B). Compared to nitrate-treated STC-1 cells, nitrite-treated cells displayed a median 1.57-fold increase in SM lipid species; similarly, compared to untreated STC-1 cells, nitrite-treated cells had a median 1.43-fold increase in SMs (Figure 5B). However, in nitrate-treated cells as compared to untreated cells, a median 1.41-fold decrease in SMs was observed. These data align with previous studies that observed elevated levels of SM in PD brains compared to controls.^52,53^ SM has also been found to be a component of Lewy inclusion bodies in the substantia nigra.^54^ Our results suggest that dysregulation of SM may be associated with intestinal PD pathogenesis as well.

A similar trend was observed for the glycerolipid, triglyceride (TG). Nitrite-treated STC-1 cells had a median 1.64-fold increase in TG lipids, as compared to nitrate-treated cells (Figure 5B); nitrite-treated cells had a median 1.46-fold increase in TG lipids when compared to untreated cells. Conversely, a median 1.65-fold decrease was observed in TG lipids in nitrate-treated cells as compared to the untreated samples (Figure 5B). While debated whether an alteration in abundance of TG lipids is linked to severe symptoms of PD, mounting evidence supports that dysregulation of TG lipid metabolism is implicated in PD pathogenesis.^55,56^ Diacylglycerol (DG), the other type of glycerolipid class identified in these data (Figure 5A), has been similarly associated with PD,^57^ and the median fold-changes of this lipid class mirror the trend observed for the TG lipids (Figure 5B). In the nitrite-treated STC-1 cells as compared to the nitrate-treated cells, a median 1.33-fold increase was displayed in DG; while in the nitrate-treated cells, a 1.46-fold decrease was observed against the untreated cells. Thus, the effect of nitrite treatment increases DG levels even more than nitrate on untreated STC-1 cells.

## DISCUSSION

Symptoms of PD are characterized by a deficit of dopamine. This occurs when the build-up of α-syn aggregates in dopaminergic neurons causes neurodegeneration and, thus, the loss of dopamine production.^58^ Levels of this critical neurotransmitter are replenished in individuals by dosing L-dopa.^59^ Unlike dopamine, L-dopa crosses the blood-brain barrier, where it is decarboxylated to form dopamine.^60^ Outside of the brain, L-dopa can also be converted to dopamine, but this is problematic because dopamine itself cannot cross the blood-brain-barrier to enter the brain and exert a therapeutic effect.^61^ To address this challenge, amino acid decarboxylase inhibitors (benserazide or carbidopa) are co-dosed with L-dopa, which prevent premature conversion of L-dopa to dopamine outside of the brain.^62^

We suspected that benserazide’s ability to limit dopamine production outside the brain (a symptomatic treatment) could simultaneously limit a putative cause of PD, namely, peripheral α-syn aggregation. Specifically, due to our findings that dopamine is a critical mediator of nitrite-induced α-syn aggregation in intestinal cells, using benserazide to limit dopamine biosynthesis was expected to limit aggregate formation. Indeed, supplying benserazide to EECs decreased the extent of pathologic α-syn aggregates formed (Figure 1B and C). These data suggest that benserazide could slow disease progression, which would be a previously unappreciated therapeutic benefit of benserazide used in conjunction with L-dopa for treatment of individuals with PD. Such strategies to limit intestinal α-syn aggregation may be particularly important because of the intestine’s capacity to serve as a reservoir for prion-like spread of aggregates. Indeed, appendix tissue is enriched in α-syn,^63^ and appendectomy significantly decreases risk of developing PD.^64,65^

Our data show that EECs are remarkably durable in the presence of α-syn aggregates. At physiologically relevant concentrations of nitrite (0.5 mM)^37^—at which a significant amount of α-syn aggregates form^2^—superoxide generation is minimal (Figure 2A), and the number of viable cells is comparable to the untreated control (Figure 2B). Only at the highest concentration of nitrite tested (50 mM) was there significant mitochondrial dysfunction and cell death. In the context of the mammalian gut, these data suggest that, despite bacterial nitrite being an oxidant, this may not drive cell-wide oxidative stress but rather have a precise impact on the cell. Theoretically, these functional intestinal cells could continuously propagate α-syn aggregates from the EECs in which they form to the enteric neurons with which they synapse, putatively advancing gut-to-brain spread of α-syn aggregates and PD pathogenesis.^7,8,66^

The absence of α-syn aggregation in nitrite-treated HeLa cells, and their consequently distinct proteome in contrast to nitrite-exposed EECs, enabled us to differentiate the relevant impacts of nitrite and α-syn aggregates on intestinal cell function. Each of the protein pathways uniquely upregulated in nitrite-treated EECs is related to the molecular mechanisms governing dopamine-dependent α-syn aggregation. Upregulation of apoptosis-induced DNA fragmentation (Figure 3D) is correlated to our findings of increased cytotoxicity and ROS at high concentrations of nitrite (Figure 2A and B). This generation of oxidative stress may similarly explain the cell’s transition to a state of cell-cycle arrest (signified by formation of SAHF) (Figure 3D). Transition to cellular senescence could also be linked to the inhibition and inactivation of the APC/C and degradation of Nek2A and Cyclin B (Figure 3D). Here, the various modes of inhibition of the APC/C could also lead to deficient tagging of α-syn aggregates for degradation, consequently leading to a build up of α-syn aggregates. Taken together, these data provide further insight into the possible spread of α-syn aggregates, which are generated by intestinal cells that remain viable and continuously forming α-syn aggregates despite accumulation of mitochondrial dysfunction and hallmarks of cell senescence.

The altered lipidomes of EECs bearing nitrite-induced α-syn aggregates display similar profiles as previous studies examining cerebrospinal fluid and the brain tissue in people with PD.^44,45^ Notably, there were dysregulated levels of sphingomyelin (SM) and triglyceride (TG) in nitrite-exposed STC-1 cells as compared to both untreated and nitrate-treated cells (Figure 5B). A similar trend was also observed for the diacylglycerol (DG) lipid species. The impact of elevated levels of TG and DG warrant further studies to examine the precise role of lipid changes on cellular function, as studies on the links between these lipids and PD severity have been contradictory.^55,56,67,68^ Curiously, a significant increase (2.55-fold) was observed for phosphatidylserine (PS), a glycerophospholipid, in the nitrite-treated STC-1 cells as compared with the untreated cells, but the P-value for PS was not significant for other comparisons (Figure 5B). Abnormalities in PS levels are linked to PD; elevated levels of PS lipids in the brains of PD patients and animal models have been recorded,^69,70^ though the mechanism by which PS is involved remains unclear.^71^ Our findings motivate further studies to examine the possible mechanistic roles of TG, DG, and PS lipids in PD pathogenesis.

Our differential lipidomic profiling provides a foundation for targeted analyses in future studies identifying molecular signatures relevant to PD etiology in the mammalian gut. Within the sulfated hexosylceramide (SHexCer), bis(monoacylglycero)phosphate (BMP), and phosphatidylcholine (PC) lipid classes, several lipid species that were upregulated in the nitrite-treated STC-1 cells as compared to the untreated cells warrant further investigations. These species include SHexCer d18:1_16:0, BMP 18:1_22:6, PC 14:0_14:0, PC 29:0, and PC 30:0 (Figure S6A). When examining the nitrite-treated STC-1 cells as compared to the nitrate-treated cells, TG 16:0_17:1_18:1 is distinguishably upregulated, while TG 14:0_26:0_22:7 is the only significantly downregulated lipid species. Notably, the differences between these two TG species is the presence of an odd-carbon chain fatty acyl. Each of the classes of lipid species have been identified in association with PD pathology in the brain;^45,46^ suggesting further investigations into the functions of these species could reveal biomarkers specific to intestinal progression of this disease.

Taken together, the studies reported herein reveal molecular and cellular pathways associated with intestinal aggregation of α-syn that is instigated by the bacterial metabolite nitrite. Collectively, our findings provide a mechanistic link between gut microbiota composition in people with PD and a putative underlying cause of the disease. Mapping this process at the level of atoms and bonds has provided insight into new therapeutic targets as well as therapies—specifically, peripheral dopamine biosynthesis and modulators of this process. We also reveal several lipid classes and species that are altered upon nitrite-induced α-syn aggregation, investigation of which may further refine mechanistic understanding of this pathologic pathway. Leveraging the mechanistic findings revealed here may lead to new avenues for improved treatment of people with PD, potentially enabling a needed strategy to stop PD pathology in the gut before it spreads to the brain.

## MATERIALS AND METHODS

### Cell Lines and Growth Conditions

Enteroendocrine STC-1 cell line (CRL-3254) and HeLa cell line (CCL-2) were obtained from the American Type Culture Collection (ATCC). Cells were cultured in Dulbecco’s Modified Eagle Medium containing 4.5 g/L glucose and 2 mM L-glutamine (DMEM, Corning) supplemented with 10% (v/v) fetal bovine serum (Life Technologies), 100 U/mL penicillin (10,000 U/mL; Gibco), 100 μg/mL streptomycin (10,000 μg/mL; Gibco), and 2.5 mg/L plasmocin prophylactic (2.5 mg/mL; Invivogen). Cells were incubated at 37 °C in a humidified atmosphere containing 5% CO_2_. Cells were serially passaged using 0.25% Trypsin-EDTA (Gibco).

### Immunofluorescence Staining

STC-1 cells were seeded onto two μ-slide 8-well high glass bottom chamber slides (Ibidi) at a density of 1 × 10^5^ viable cells/well and incubated for 48 h. Growth medium was then replaced with fresh DMEM supplemented with powdered sodium nitrate (50 mM; ThermoScientific) or sodium nitrite (50 mM; Sigma), and with benserazide (0.01 mM; Cayman Chemical), or vehicle and then filter-sterilized. Cells were incubated in this appended DMEM for another 24 h before fixation. Fixation was performed at room temperature (RT) using 10% formalin (Fisher Scientific) for 20 min. Cells were then washed twice for 5 min using sterile-filtered phosphate-buffered saline (PBS, pH 7.4). All wash steps used PBS for 5 min each and were performed at RT. Cells were then permeabilized using 0.1% Triton X-100 (Bio-Rad) in PBS for 20 min at RT, and subsequently were washed twice. The cells were then blocked for 1 h at RT using 5% normal goat serum (Thermo Scientific) and 0.2% bovine serum albumin (BSA; Fisher Scientific) in PBS. Thereafter, the samples were washed three times, then incubated in PBS containing 0.2% BSA with either MJFR-14 anti-aggregate primary antibody (Abcam, ab209538; 1:250 dilution), isotype control (Abcam, ab172730; 1:566 dilution), or vehicle overnight at 4 °C. After washing three times, all chambers were incubated with anti-goat Alexa Fluor-488 secondary antibody (Abcam, ab150077;1:500 dilution) for 1 h in the dark at RT. All subsequent steps were performed in the dark. After washing three times, ∼10 drops of VECTASHEILD PLUS Antifade Mounting Medium with DAPI (Vector Laboratories) were added to each well. The chamber slides were stored at 4 °C in the dark until microscopy images were gathered.

### Structured Illumination Microscopy

A Zeiss Elyra 7 super-resolution microscope with a 63x oil immersion lens was used for acquiring three images for each sample. Images were collected using 405 and 488 nm laser lines for excitation. Emission filters BP 420−480 nm and BP 495−550 nm were selected. Z-stack images were obtained and processed using SIM^2^ scaled to the raw image. Quantification of the fluorescence signal for each sample was determined by obtaining the mean intensity of the maximum intensity projection for each image given by Zen Black 3.0 software. The mean intensity per cell was then calculated. To account for any background fluorescence signal, the mean intensity per cell for the respective untreated sample was subtracted for each independent replicate. Results are expressed as arbitrary units (au) of the background-subtracted mean intensity per cell. GraphPad Prism (San Diego, CA) was utilized for graphing, and statistical analyses using one-way ANOVA were performed to compare samples.

### Mitochondrial Superoxide (MitoSOX) Detection

Mitochondrial dysfunction was investigated as a function of superoxide generation using a MitoSOX fluorescence-based assay kit (Abcam, ab219943). STC-1 cells were seeded onto a 96-well clear-bottom black-wall plate (Greiner) at a density of 6.5 x 10^4^ viable cells/well and incubated for 24 h. Growth medium was aspirated and cells were washed once with PBS. FluoroBrite DMEM (Gibco) supplemented with powdered sodium nitrite (0.005-50 mM), sodium nitrate (0.005-50 mM), antimycin A (50 μM; Enzo Life Sciences), or vehicle was sterile-filtered and then supplied. Each condition was performed in triplicate. Cells were incubated for another 24 h, then stained with 100 μL MitoROS 580 working solution (prepared as specified by the manufacturer) for 1 h in the dark. Changes in fluorescence intensity were monitored with a BioTek plate reader (Ex/Em = 540/590 nm). Data were plotted using Prism.

### MTT Cell Viability Assay

STC-1 cells were seeded onto a 96-well clear plate (Greiner) at a density of 6.5 x 10^4^ viable cells/well and incubated for 24 h. Growth medium was then replaced with fresh DMEM supplemented with either sodium nitrite (0.005-50 mM), sodium nitrate (0.005-50 mM), or vehicle and sterile-filtered. Each condition was tested in triplicate using supplemented DMEM, with cells incubated for another 24 h. Treated growth medium was then aspirated and wells were washed twice with PBS. Cells were incubated with 100 μL sterile-filtered MTT working solution (0.5 mg/mL thiazolyl blue tetrazolium bromide in PBS, Thermo Fisher Scientific) for 4 h. Once the MTT solution was carefully removed, cells were lysed using 100 μL DMSO (Sigma Aldrich). A BioTek plate reader was used to measure the absorbance at 570 nm to quantify the formazan dye product. The reported live cell percentage was calculated for each well by assuming the untreated cells were 100% viable. The corresponding mean background absorbance from the DMSO-only wells was subtracted for each independent experiment. Data were plotted using Prism.

### Stimulating Aggregation for Proteomics Analysis

STC-1 cells were seeded into T-75 flasks. After incubating for 48 h under standard growth conditions described above, the growth medium was replaced with fresh, sterile-filtered media supplemented with either sodium nitrite (0.005-50 mM), sodium nitrate (0.005-50 mM), or vehicle. Cells were then incubated for another 24 h before lifting with 0.25% Trypsin-EDTA. After neutralizing, samples were centrifuged at 125 x g for 5 min. Media was aspirated, and cells were resuspended in PBS. Cells were then washed three additional times with fresh PBS, centrifuging at 500 x g for 5 min after each resuspension before aspirating. Pellets were stored at -80 °C until thawing for proteomics analysis.

### Protein Digestion and Liquid Chromatography-Mass Spectrometry (LC-MS) Analysis for Proteomics

Mouse-derived STC-1 and HeLa cells were cultured and treated with 50 mM nitrate, 50 mM nitrite, and vehicle control as described above. To each pelleted sample, 100 μL of lysis buffer [5% sodium dodecyl sulfate (SDS) and 50 mM triethylammonium bicarbonate buffer (TEAB)] supplemented with Pierce Protease Inhibitor cocktail (Thermo Fisher Scientific, Waltham, MA) and phosphatase inhibitors (10 mM sodium pyrophosphate, 1 mM phenylmethyl sulfonyl fluoride, 1 mM sodium orthovanadate, 1 mM sodium fluoride, and 1 mM *β*-glycerol phosphate) was used. After 10 min of incubation in the lysis buffer on ice, samples were sonicated for 3 cycles of 10 pulses at 30% power using a QSonica probe sonicator. Samples were then centrifuged at 14,000 x g for 10 min at 4 °C. The supernatant from each sample was transferred to a new microcentrifuge tube, and the bicinchoninic acid assay (BCA) (Thermo Fisher Scientific, Waltham, MA) was performed to determine protein concentration. In total, 100 μg of protein from each sample was taken for protein digestion using the S-Trap Micro Spin Column Digestion protocol (Protifi, Huntington, NY) with minor changes per the manufacturer’s protocol. Briefly, dithiothreitol was added to a final concentration of 20 mM, and samples were heated to 95°C for 10 min. After cooling to room temperature, samples were alkylated with 40 mM iodoacetamide and incubated for 30 min in the dark. Samples were then acidified with 1.2% (v/v) phosphoric acid (final concentration), followed by brief vortexing. Next, 150 μL of S-Trap binding buffer [90% (v/v) methanol and 100 mM TEAB] was added, and samples were mixed once again. The mixtures were then loaded onto the spin column and centrifuged at 4000 x g for 30 sec. Washing steps with the binding buffer were repeated five times in total, prior to the addition of trypsin [1:25 enzyme (μg):protein (μg) ratio] in 40 μL of 50 mM TEAB. Digestion was carried out by incubation at 37 °C overnight. Peptides were eluted with 40 μL of 50 mM TEAB, 40 μL of 0.2% formic acid (FA), and 40 μL of 50% acetonitrile (ACN), each followed by a 1 min centrifugation at 4000 x g. Each proteolytic peptide sample was dried *in vacuo* and resuspended in 100 μL of 0.1% formic acid prior to analysis.

Samples were injected (0.5 μL) and analyzed using the label-free quantification (LFQ) approach on a Q-Exactive mass spectrometer (Thermo Fisher Scientific, Waltham, MA) coupled with an Agilent 1260 Infinity nanoLC system (Agilent Technologies, Santa Clara, CA). Proteolytic peptides were loaded onto a Thermo NanoViper trap column (75 μm × 2 cm, 3 μm C18, 100 Å) (Thermo Fisher Scientific, Waltham, MA). After 10 min of washing with 0.1% FA at 2 μL/min, peptides were separated using an Agilent Zorbax 300SB-C18 analytical column (0.075 × 150 mm, 3.5 μm, 300 Å) (Agilent, Santa Clara, CA) with a 120 min gradient, from 5 to 60% ACN with 0.1% FA, at a flow rate of 0.25 μL/min. Data collection was carried out using data-dependent acquisition (DDA). Peptides were ionized at a spray voltage of 1.8 kV, and the capillary temperature was set at 250 °C. A full MS1 scan was performed from 375 to 1600 m/z at 70,000 resolution. The automatic gain control (AGC) target was set at 1 × 10^6^ ions, and the maximum injection time (IT) was set to 100 ms. The ten most abundant peaks within a MS1 spectrum were isolated for MS2, with an isolation width at 1.5 m/z and dynamic active exclusion was set for 20 s. MS2 spectra were collected at 17,500 resolution, for a maximum of 50 ms or a minimum of 1×10^5^ ions. Normalized collision energy (NCE) for HCD fragmentation was set at 27%. Ions observed with charges of +1 and greater than +6 were excluded from MS2 analysis.

Database searching was carried out using Proteome Discoverer 3.1 software (Thermo Fisher Scientific, Waltham, MA), and searched against the UniProt *Mus musculus* database (v2023-06-28). The precursor assignment used peptides with masses from 350 to 9600 Da, cleaved by trypsin, containing 7 to 30 amino acids with 2 or fewer missed cleavages. Precursor ion peak intensities were used for quantification with a mass error set at 10 ppm and fragment masses were searched with a tolerance of 0.02 Da. Carbamidomethyl (C, +57.021 Da) was set as a static modification, and acetylation (protein N-terminus, +42.011 Da) was set as a dynamic modification. Filtering was then applied such that only high-confidence proteins and peptides (FDR ≤0.01) remained. All proteins were required to have at least 2 unique peptide matches. No quantitative value correction was applied. CHIMERYS intelligent search algorithm was included in all database searches. The protein abundance was calculated by summing the abundance of all unique and razor peptides normalized to the total peptide amount, and protein ratios were derived from the pairwise peptide group ratios. No imputation was executed for missing values. The maximum or minimum allowed fold-change and fold-change of proteins with missing values across all biological replicates within a condition are 100 and 0.01. A background-based *t*-test on the normalized protein abundance was conducted in Proteome Discoverer to determine significant changes for all proteins detected including α-syn. Within the differential expression results, we considered proteins with an adjusted P value < 0.05 as statistically significant. Overlap analysis of the treatment groups was performed using Venny 2.1.0 (https://bioinfogp.cnb.csic.es/tools/venny/). Differential proteins were subject to pathway analysis using the Reactome database (https://reactome.org/PathwayBrowser/#TOOL=AT).

### Lipid Extraction and LC-MS Analysis for Lipidomics

Mouse-derived STC-1 cells were cultured and treated with 50 mM nitrate, 50 mM nitrite, or vehicle control as described above. The cell pellets were homogenized on ice-cold lysis buffer (1× PBS containing protease and phosphatase inhibitors, as described above). Protein concentration was measured using the Pierce BCA Protein Assay (Thermo Fisher Scientific) according to the manufacturer’s protocol. Organic extraction of lipids was performed using the Folch lipid extraction method from a 50 μg protein equivalent of the lysates.^72^ Here, 5 μL of SPLASH Lipidomix was added as an internal standard prior to extraction. The organic layer resulting from the lipid extraction was dried at 30 °C *in vacuo* and resuspended in 50 μL methanol-chloroform (3:1, v/v) prior to analysis.

LC-MS analysis of the lipid extracts was performed using an Agilent 6550 iFunnel Q-TOF mass spectrometer coupled with Agilent 1260 infinity HPLC. Pooled samples were prepared by combining 5 μL of each sample and an iterative MS/MS workflow was performed in the Mass Hunter acquisition software across seven injections of the pooled samples to generate a library for analysis of individual samples. The chromatographic separation included mobile phase A of H_2_O:methanol 90:10 (v/v) with ammonium acetate (5 mM) and B of isopropanol:methanol:acetonitrile 5:3:2 (v/v/v) with ammonium acetate (5 mM). For both ion modes, 3 μL of the resuspended lipid extracts were loaded onto a 2.1 × 100 mm Agilent Poroshell C18, 2.7 μm column (Agilent Technologies Inc., Santa Clara, CA, USA) and separation was performed using the following gradient: 70% B at 0– 1 min, 86% B at 3.5–10 min, 100% B at 11–17 min and at a flow rate of 300 μL min^−1^. A post-column equilibration time of 10 min was used for all runs. The mass spectrometer was operated in 2 GHz extended dynamic range mode employing precursor ion analysis for relative quantification experiments in both negative and positive ion modes. In the negative-ion mode, internal reference mass calibration ions m/z 119.0363 and 980.0163 were used. In positive-ion mode, internal reference mass calibration ions m/z 121.0509 and 922.0098 were used. Source parameters were as follows: gas temp, 200 °C; gas flow, 11 L min^-1^; nebulizer, 35 psi; sheath gas temp, 250 °C; sheath gas flow, 12 L min^-1^; VCap, 3500 V; and fragmentor, 145 V. MS1 acquisition mode was applied, m/z range was set to 150–1700 with a scan rate of 4.00 spectra per second.

Lipids assignments were made based on fragmentation matching (MS/MS) to the LipidBlast library using the Lipid Annotator software (Agilent Technologies Inc., Santa Clara, CA, USA). Retention time for each lipid was aligned to ±0.1 min using a mass accuracy window of < 5 ppm and extracted-ion peaks integrated using the Agilent integrator in the Profinder software. Each of the integrated peaks was manually reviewed for retention time and fragmentation matching. The processed data file was exported and imported into the Mass Profiler Professional software (Agilent Technologies Inc., Santa Clara, CA, USA) where each data set was analyzed separately in positive- and negative-ion modes. Additionally, a lipid class-based normalization workflow was performed in the Mass Profiler Professional software to account for any variation in extraction efficiency across samples. In both ion modes, all lipid intensities were baselined to median intensity. Lipids that were not present in all biological replicates of either condition were further filtered. Next, a list of altered lipids for each ion mode was generated based on individual lipid intensities, then one-way ANOVA (P < 0.05) was used to determine significant alterations. Lipids identified in both negative and positive mode are reported based on the mode with higher statistical significance (lower P value).

## ASSOCIATED CONTENT

All raw mass spectrometry data was deposited and is available at the MassIVE data (http://massive.ucsd.edu) repository (MSV000098917) with password of 0SWGDfQX4ZmG707c.

## Supplementary Materials

The Supporting Information contains Figures S1–S6 and Tables S1-S5.

- Figure S1: Representative images of fixed STC-1 cells incubated for 24 h with NO_3_^−^, NO_2_^−^, and/or Benz, or vehicle, probed with isotype control.
- Figure S2: Representative images of fixed STC-1s confirming aggregation was induced by nitrite exposure before lysis.
- Figure S3: Volcano plot from proteomic analysis of STC-1 enteroendocrine cells when exposed to 50 mM nitrate as compared to untreated controls.
- Figure S4: Representative images of fixed HeLa cells incubated for 24 h with NO_3_^−^, NO_2_^−^, or vehicle, probed with primary antibody MJFR-14 or isotype control.
- Figure S5: Volcano plot from proteomic analysis of HeLa cells when exposed to 50 mM nitrate as compared to untreated control cells.
- Figure S6: Volcano plots from untargeted lipidomic analysis of STC-1 enteroendocrine cells when exposed to nitrite, nitrate, or vehicle.
- Table S1: Complete proteomic analysis of STC-1 enteroendocrine cells.
- Table S2: Proteins uniquely altered in nitrite-treated STC-1 enteroendocrine cells.
- Table S3: Complete proteomic analysis of HeLa cells.
- Table S4: Proteins uniquely altered in nitrite-treated HeLa cells.
- Table S5: Complete lipidomic analysis of STC-1 enteroendocrine cells.
- Table S6: Abbreviations of lipids identified.

## AUTHOR INFORMATION

## Author Contributions

J.M.B., Y.Y., and D.T. performed experiments. J.M.B., Y.Y., D.T., S.M.C., and E.N.B designed experiments and conducted data analysis. J.M.B., Y.Y., S.M.C., and E.N.B. wrote and edited the manuscript. S.M.C. and E.N.B. acquired funds and provided project supervision and administration. All authors read and approved the final version of the manuscript.

## Notes

The authors declare no conflicts of interest. All authors read and approved the final version of the manuscript. Data generated or analyzed during this study are included in the manuscript, and supporting files are available from the corresponding author upon reasonable request.

## ACKNOWLEDGMENTS

Financial support for this publication was provided by Scialog grants #28647 and #28648, sponsored jointly by Research Corporation for Science Advancement, the Frederick Gardner Cottrell Foundation, and the Paul G. Allen Frontiers Group. Research reported in this publication was also supported by the National Institute of Neurological Disorders and Stroke of the National Institutes of Health under Award Number R01NS138879. The content is solely the responsibility of the authors and does not necessarily represent the official views of the National Institutes of Health. SMC was also supported by the Walder Foundation. This study was made possible in part through access to the Optical Biology Core Facility of the Developmental Biology Center, a shared resource at the University of California, Irvine. The table of contents graphic was created with BioRender.com (agreement number CS28OJFJSK).

## Abbreviations

PD: Parkinson’s disease
α-syn: α-synuclein
EECs: enteroendocrine cells
ROS: reactive oxygen species
AMA: antimycin A
MitoSOX: mitochondrial superoxide
MTT: thiazolyl blue tetrazolium bromide
SAHF: senescence-associated heterochromatin foci
HMOX1: heme oxygenase-1
APC/C: anaphase-promoting complex/cyclosome
LC–MS: liquid chromatography-mass spectrometry
BMP: bis(monoacylglycero)phosphate
CL: cardiolipin
etherPC: ether-linked phosphatidylcholine
etherPE: ether-linked phosphatidylethanolamine
LPC: lysophosphatidylcholine
LPE: lysophosphatidylethanolamine
PC: phosphatidylcholines
PE: phosphatidylethanolamines
PG: phosphatidylglycerol
PI: phosphatidylinositol
PS: phosphatidylserine
Cer_ADS: α-hydroxy-dihydrosphingosine ceramide
Cer_NDS: non-hydroxy-dihydrosphingosine ceramide
Cer_NS: non-hydroxy-sphingosine ceramide
CerP: ceramide phosphate
GM3: monosialodihexosylganglioside
HexCer_NS: hexosylceramide containing non-hydroxy fatty acid and sphingosine
SHexCer: sulfated hexosylceramide
SM: sphingomyelin
DG: diacylglycerol
TG: triglycerides
ACar: acylcarnitine
FA: fatty acyl
FAHFA: fatty acyl ester of hydroxy fatty acid
CE: cholesterol ester

