## Supplemental Figures for "Multiomic Analysis Reveals Molecular Pathways Associated with Intestinal Aggregation of α-Synuclein"

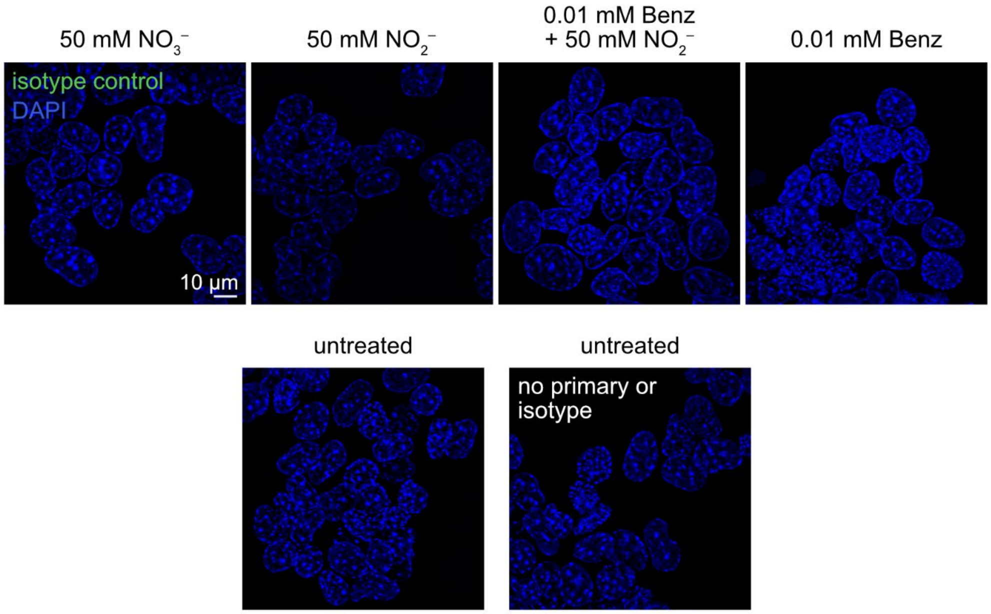


**Figure S1. Nitrite-induced dopamine-dependent α-syn aggregation in enteroendocrine cells (EECs) is reduced in the presence of benserazide.** Representative images of fixed STC-1 cells incubated for 24 h with nitrate, nitrite, and/or benserazide (Benz) probed with isotype control (blue = DAPI-stained nuclei, green = non-specific interactions).

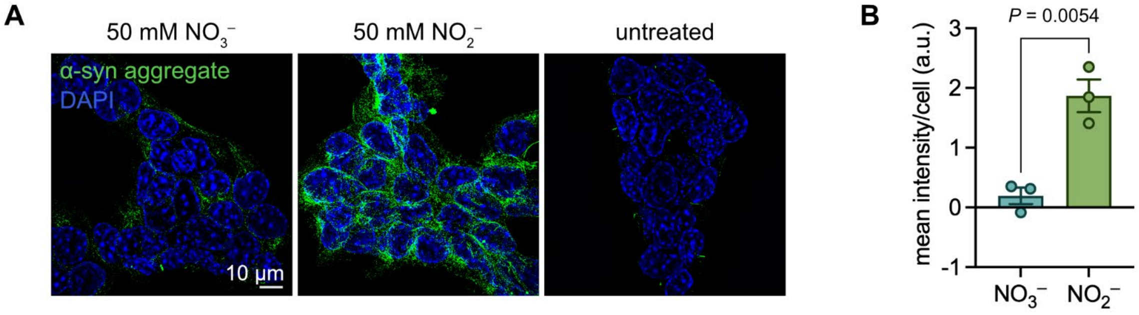


**Figure S2. α-Syn aggregation in enteroendocrine cells (EECs) was induced by nitrite exposure prior to cell lysis for proteomic and lipidomic analysis.** (A) Representative images of fixed STC-1 cells incubated for 24 h with nitrate, nitrite, or vehicle probed with anti-aggregate primary antibody MJFR-14 (blue = DAPI-stained nuclei, green = α-syn aggregation). (B) Mean fluorescence intensity per cell was quantified from maximum intensity projections acquired by structured illumination microscopy ($n$ = 3 technical replicates for each, bars denote mean ± S.E.M.; significance determined by unpaired t-test).

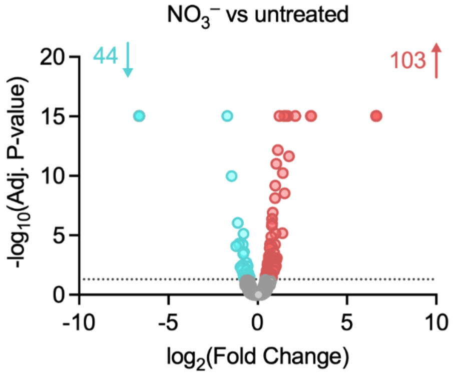


**Figure S3. Proteomic analysis of STC-1 enteroendocrine cells (EECs) when exposed to 50 mM nitrate as compared to untreated controls.** Volcano plot showing differentially expressed proteins (cyan = downregulated, coral = upregulated). The P-value cutoff was 0.05. In the nitrate group, 3704 proteins were identified; 44 were downregulated and 103 were upregulated as compared with untreated controls.

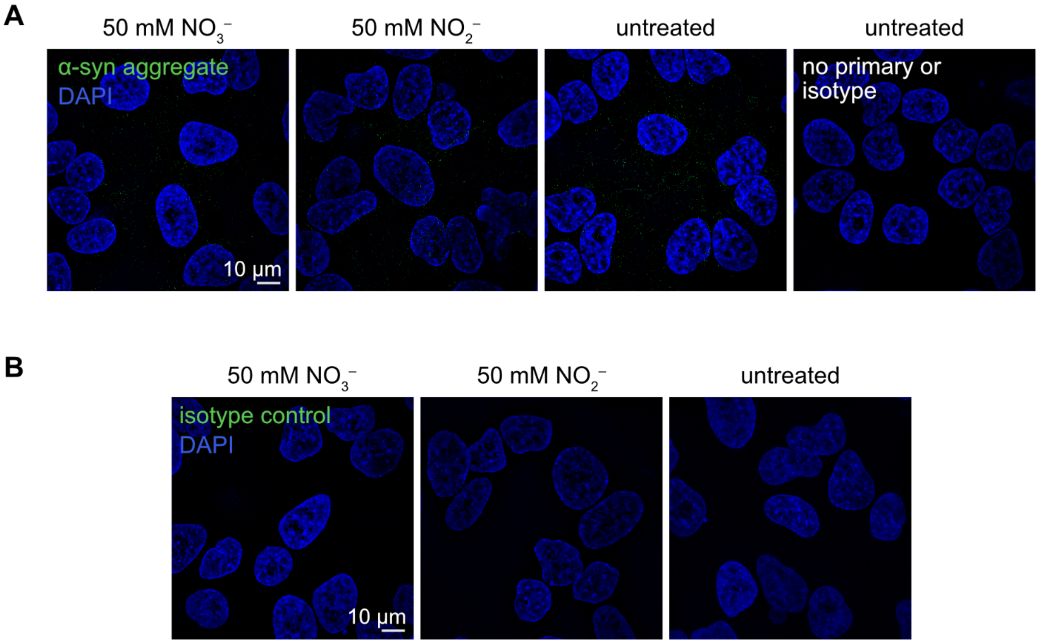


**Figure S4. Supplying 50 mM nitrite to HeLa cells does not induce aggregation of natively expressed α-syn.** Representative images of fixed HeLa cells incubated for 24 h with nitrate, nitrite, or vehicle probed with (A) anti-aggregate primary antibody MJFR-14 (blue = DAPI-stained nuclei, green = α-syn aggregation) or (B) isotype control (blue = DAPI-stained nuclei, green = non-specific interactions).

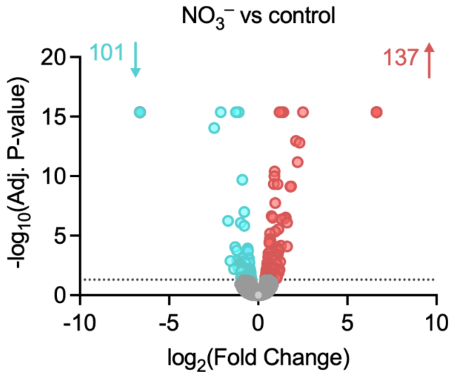


**Figure S5. Proteomic analysis of dopamine-free HeLa cells when exposed to 50 mM nitrate as compared with untreated controls.** Volcano plot showing differentially expressed proteins (cyan = downregulated, coral = upregulated). The P-value cutoff was 0.05. In the nitrate group, 4031 proteins were identified; 101 were downregulated and 137 were upregulated as compared with untreated controls.

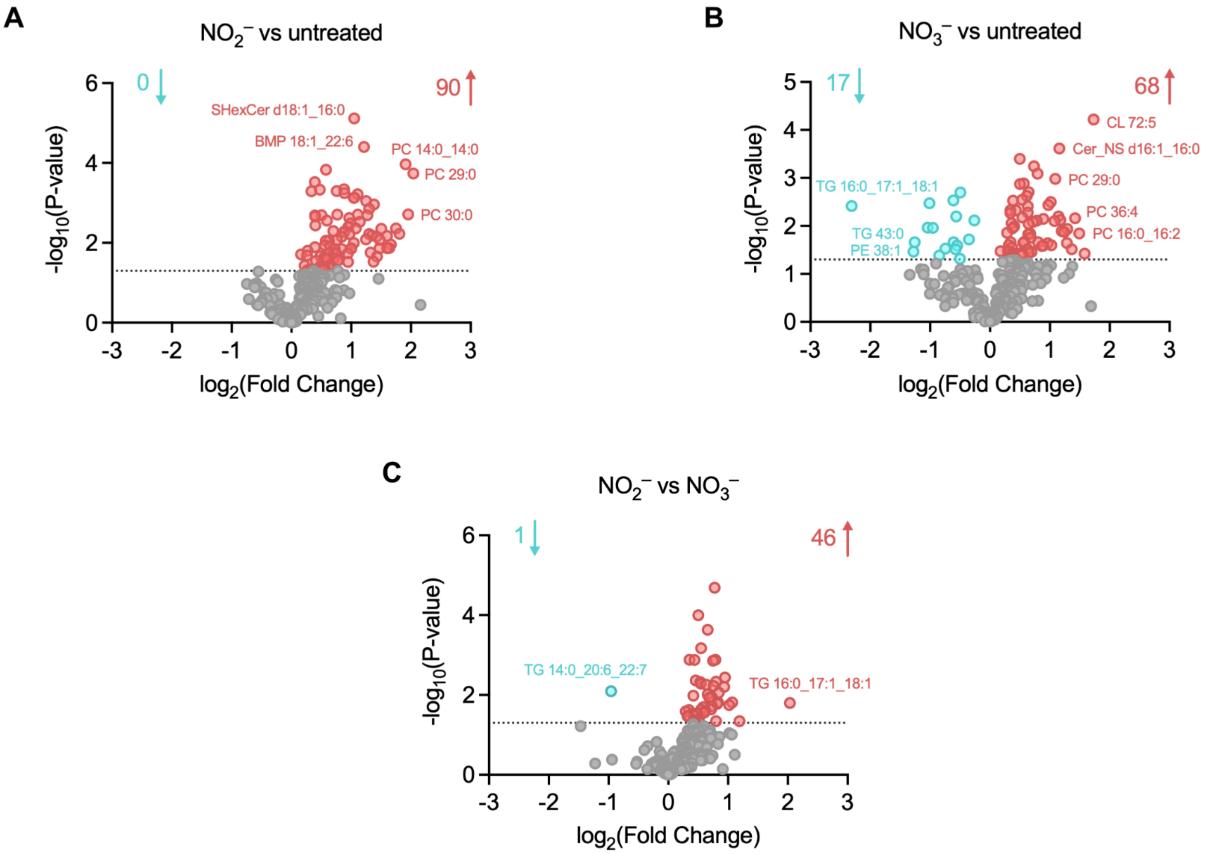


**Figure S6. Lipidomic analysis of STC-1 enteroendocrine cells (EECs) when exposed to nitrite, nitrate, or vehicle.** Volcano plots showing lipids with differentially altered abundance (cyan = downregulated, coral = upregulated) in STC-1 cells. The P-value cutoff was 0.05. In positive and negative mode, a total of 282 lipids were identified. (A) In STC-1 cells exposed to 50 mM nitrite compared to untreated cells, 0 lipids were downregulated and 90 were upregulated. (B) In the 50 mM nitrate-treated STC-1 cells, 17 lipids were downregulated and 68 were upregulated as compared with untreated cells. (C) Compared to STC-1 cells exposed to 50 mM nitrate, 1 lipid species was downregulated and 46 were upregulated in those cells treated with 50 mM nitrite.

**Table S6. Abbreviations of lipid classes identified.**

| | |
|---|---|
| BMP | bis(monoacylglycero)phosphate |
| CL | cardiolipin |
| etherPC | ether-linked phosphatidylcholine |
| etherPE | ether-linked phosphatidylethanolamine |
| LPC | lysophosphatidylcholine |
| LPE | lysophosphatidylethanolamine |
| PC | phosphatidylcholines |
| PE | phosphatidylethanolamines |
| PG | phosphatidylglycerol |
| PI | phosphatidylinositol |
| PS | phosphatidylserine |
| Cer_ADS | α-hydroxy-dihydrosphingosine ceramide |
| Cer_NDS | non-hydroxy-dihydrosphingosine ceramide |

| | |
|---|---|
| Cer_NS | non-hydroxy-sphingosine ceramide |
| CerP | ceramide phosphate |
| GM3 | monosialodihexosylganglioside |
| HexCer_NS | hexosylceramide containing non-hydroxy fatty acid and sphingosine |
| SHexCer | sulfated hexosylceramide |
| SM | sphingomyelin |
| DG | diacylglycerol |
| TG | triglycerides |
| ACar | acylcarnitine |
| FA | fatty acyl |
| FAHFA | fatty acyl ester of hydroxy fatty acid |
| CE | cholesterol ester |
